# Copy Number Variants and heritability estimates on UKBiobank data

**DOI:** 10.1101/2023.07.24.550287

**Authors:** Jawad Boulahfa, Edith Le Floch, Morgane Pierre-Jean, Jean-François Deleuze, Claire Dandine-Roulland

## Abstract

Copy Number Variants (CNVs) are sometimes used to perform association studies. The aim of this paper was to study the use of CNVs in another context: heritability estimation. We wanted to assess the impact of using CNVs in these estimates, either alone, or in conjunction with Single Nucleotide Polymorphisms (SNPs). Using real SNP and CNV data from UK Biobank, we simulated phenotypes depending either on one or the two type(s) of data. We showed that mixed models, usually used for estimating heritability on SNP data, were also capable of estimating CNV heritability and to properly decipher between CNV and SNP heritabilities when phenotypes depend on both types of data. However CNV heritability estimation becomes more challenging when it is only supported by the few relatively common CNVs. Finally we estimated CNV and SNP heritabilities for two real phenotypes from UK Biobank (height and hypertension) but only hypertension showed a small but non-null CNV heritability of about 1.7%.

## Introduction

Copy Number Variants (CNVs) are sometimes used to perform association studies, as can be seen in the articles (1), (2) and (3) and in UKBioBank data (4). The aim of this paper was to study the use of CNVs in another context: heritability estimation. We wanted to assess the impact of using CNVs in these estimates, either alone, or in conjunction with Single Nucleotide Polymorphisms (SNPs).

For our research, we used UKBioBank data which include genetic and phenotypic information on around 500 000 individuals recruited in the United Kingdom. More precisely, we considered an homogeneous subsample of 10 000 individuals and computed heritabilities of CNVs for simulated phenotypes, height measurements and hypertension diagnostics.

## Materials

### UKBiobank dataset

The UKBiobank project began in 2006, with the aim of collecting and tracking genetic and medical data on around half a million individuals in Great Britain, in order to advance health research (5). The UKBiobank dataset contains genetic and phenotypic information on around 500 000 individuals recruited in the United Kingdom between 2006 and 2010. We can notify that the vast majority of individuals are of British origin (see Supplementary Fig. 1). Phenotypic information include the results of verbal interviews and on-site questionnaires, as well as measurements of anthropometric traits (height, BMI, etc.) obtained during the visit to one of the 22 UKBiobank project assessment centers. Some individuals’ information has been filled in more than once (as they were invited to make several visits), and in this case we have access to data from each of these assessments. Here, we only considered information from the first visit, as this is common to all individuals. In this paper, we will only analyse as phenotype simulation data and height and hypertension phenotypes.

### Genotypic data

Genotypic data of the UKBioBank individuals are detailed in the article (6) as well as the ressource 155580. Here, you only used cleaned SNPs Affymetrix genotypic array data (784 256 autosomal SNPs). Principal component analysis (PCA) was performed by consortium on a subset of 407 219 unrelated individuals and 147 604 SNPs in low linkage disequilibrium (6) and unused individuals were projected onto the resulting principal components (PCs) to obtain scores for the entire cohort (see Supplementary Fig. 2 and 3). This PCs will be used for correction of population stratification in our models.

In order to limit computation times, we extracted a subsample of 10 000 individuals from UKBioBank dataset. To do this, we began by selecting only individuals of British origin and Caucasian genetic group. Next, we calculated the mean of the first ten principal components and selected the 12 000 individuals closest to this mean in the sense of the Euclidean norm. Finally, we randomly selected 10 000 individuals from among these. This approach allows us to take into account possible population stratification, while introducing variability into the construction of our sub-sample. It’s important to note that all the batches as well as the two chips mentioned of the UKBioBank dataset are present in our subsample. We have therefore taken care to include this information as a fixed effect in our models.

#### Quality controls on SNPs

Initially, our SNP data comprise 10 000 individuals and 784 256 autosomal SNPs. Here we describe the quality checks that had to be carried out on these data. First of all, we removed monomorphic and non-biallelic SNPs. Then, we only kept individuals with a *callrate* greater than or equal to 0.95 on selected SNPs. We also decided to keep only SNPs with a *callrate* greater than or equal to 0.99 and no monomorphic. The Hardy-Weinberg disequilibrium test was applied and we removed SNPs with a p-value less than or equal to 10^*−*7^. We also removed individuals with extreme heterozygotie value or hight proportion of uninformed SNPs flagged by Consortium (6).

We also decide to removed related individuals. For that, we computed GRM on common (*MAF ≤*0.01) SNPs in low linkage disequilibrium (*r*^2^ ≥0.2). Then, we removed one individual at random from each pair where the genetic correlation coefficient is greater than or equal to 0.025 (7). In this step, 451 related paires are detected.

### CNVs data

In additions to SNPs, CNVs were extracted from SNP data using the **PennCNV** tool (8) and the pipeline described in (3). During this extraction, 2 871 individuals have no CNVs detected for them and 235 005 CNVs were called. After calling, the same CNV may have a slightly different start and/or end position(s) depending on the individual. Then, referring to the article (8), we simplify our calculations by assuming that each CNV has the same positions for all individuals and we computed a CNV count matrix by SNP position. We obtained 421 640 positions with one CNV. We can note that there are no Loss of Heterozygosity (LOH) case our CNVs data, so the 2 copies case in this count matrix indicate a normal copy number for the SNP in question (see Supplementary Note 1).

#### Quality controls on CNVs

In the count matrix, we can note the presence of positions without a given identifier or duplicate positions. We decided to remove these positions. We also removed all positions with the same number of copies for all individuals.

## Methods

### Heritability estimation

Heritability in the broad sense, noted *H*^2^, is the proportion of phenotypic variance explained by genetics. The effects of genetics can be summarized in three components: additive effects (effects linked to the alleles of a variant), dominant effects (effects linked to interactions between the alleles of a variant), and epistasis effects (effects linked to interactions between variants). In this paper, we only considered the restricted heritability, denoted *h*^2^ which is the proportion of phenotypic variance explained by the additive effects of variants. Here, we will refer to restricted heritability as heritability.

#### Mixed Linear Model

We estimate heritability using linear mixed model (9) which includes the so-called fixed effects (derived from features such as age, for example), from the so-called random effects (corresponding to the additive genetic effects of SNPs and/or CNVs). The linear mixed model, in the genetic context, is written as:

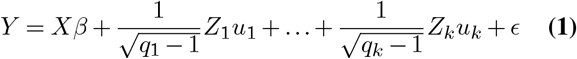

with *Y* the *n* size vector phenotype of interest (continuous quantitative) with *n* the number of individuals, *X* the feature matrix of size *n × p*) (including a column of 1 for the intercept), *β* the fixed effects *p* vector, 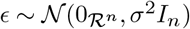 the environmental/error effects vector with *σ*^2^ ∈ *R*_+_ and, for all *i* ϵ ⟦ 1, *k* ⟧, *q*_*i*_ the number of markers in marker set *i, Z* _*i*_ the matrix of centered and reduced markers of size *n × q*_*i*_ and 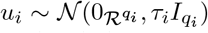, the vector of random effects (associated with the set of markers *i*) with *τ*_*i*_ ∈ *R*_+_.

The linear mixed model can also be written as:

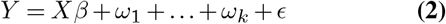

with 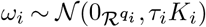 and

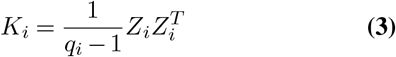

of size *n ×n*.

In case of SNPs, *K*_*i*_ is the Genetic Relatedness Matrix (GRM) (9) which is an estimator of the variance-covariance matrix of individuals calculated on markers.

#### Heritability estimates

Parameter estimators, 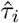 and 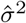, were be computed by AIREML(10) in Eq. (2). Then, the heritability 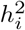 of the markers in set *i* is estimated by:

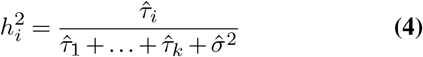

#### Used models

In our particular case, we consider two sets of markers, the first with all SNPs and the second one for CNV counts. Then, the complete linear mixed model is written as:

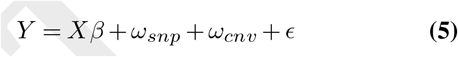

with 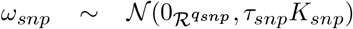, 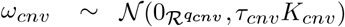, *K*_*snp*_ the GRM computed on the genotype matrix of standardized SNPs and *K*_*cnv*_ an estimator of the variance-covariance matrix of the individuals calculated on the copy numbers of the CNVs. The feature matrix *X* included intercept, genomic sex, age, the batch number, the sequencing chip (Axiom or BiLEVE) and possibly first PCs computed on 10 000 subsamples (various cases have been considered).

We also considered linear mixed models with two random effect vectors for common and rare CNVs (see Results for details).

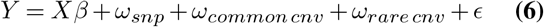

with previous notations and *ω*_*commoncnv*_ *∼ 𝒩* (0_*R qcommon cnv*_, *τ*_*common cnv*_ *K*_*common cnv*_), *ω*_*rare cnv*_ *∼ 𝒩* (0_*R qrare cnv*_, *τ*_*rare cnv*_ *K*_*rare cnv*_), *K*_*common cnv*_ and *K*_*rarecnv*_ estimators of the variance-covariance matrix of the individuals calculated on the copy numbers of the common and rare CNVs respectively.

The linear mixed models with only one random effect have also been estimated for SNPs

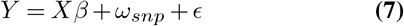

or CNVs (all, common or rare)

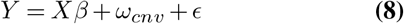

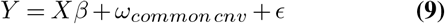

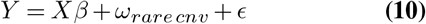

with the same notations.

#### Binary phenotype

In this paper, we also analysed a binary phenotype which define a particular disease. For that, the Gaussian linear mixed models given in previous paragraph can be used if they are followed by a linear transformation of this estimate to obtain the heritability on the liability scale (11).

### Phenotype Simulations

For simulations, we have only considered the cleaned individuals. We simulated quantitative phenotypes under linear mixed models of Eq. (5) and Eq. (6). We considered several scenarios:

1. the phenotype depends on SNPs

2. the phenotype depends on CNVs

3. the phenotype depends on common CNVs

4. the phenotype depends on rare CNVs

5. the phenotype depends on SNPs and CNVs

6. the phenotype depends on SNPs and common CNVs

7. the phenotype depends on SNPs and rare CNVs

For each scenario, many PC numbers included in features and heritability values have been tested for simulation and the simulation process was repeated 100 times for each parameter combination. The linear mixed models have then been estimated on these sets of 100 simulations. We can note that, in each case, the PCs were not included in feature matrix for simulation because add them did not change conclusions (results not shown).

Results for scenarios (3), (4), (6) and (7) are shown in Supplemetary Note 2.

### Resampling for height and hypertension phenotype

Concerning heritability estimates on observed height or hypertension vector, we tested two different approaches:

1. estimating point wise the heritability of height, as a function of the number (⟦ 0, 20⟧) of principal components we include in fixed effects. The first approach consists of building a single model, and therefore producing a single estimate, for each choice of number of principal components to be included as fixed effects.

2. estimating the heritability of height as a function of the number ({0, 5, 10, 15, 20}) of principal components that we include in fixed effects, after resampling individuals (100 resamples of 5 000 individuals). The second approach is to build several models for each choice of number of principal components, each model using a different subsample of our 7 207 individuals. This produces several estimates for each choice of number of principal components. We can then assess the variability of these estimates. In order to reduce computation times, the GRM and the estimator of the variance-covariance matrix of the individuals calculated on the copy numbers of the CNVs are not recalculated for each new sample, but derived from the initial truncated matrices (only the selected individuals are kept).

### Software

All analyses and computations were performed with package R gaston [1.5.7] (12).

## Results

### Quality controls

#### Individuals filtering

To summarize, individuals are removed from our 10 000 individual-subsample after quality controls on the SNPs and CNVs data as soon as at least one of these criteria was met:

- *callrate* strictly less than 0.95;
- presence of NA’s for anthropometric traits;
- *outliers* for heterozygosity or the proportion of uninformed SNPs (see Materials);
- genetic correlation coefficient greater than or equal to 0.025;
- number of CNVs detected greater than or equal to 30. In the end, we removed 2 793 individuals.

PCA was also applied on our cleaned and pruned data (see Method). We can see that our subsample is homogeneous and so the population stratification is limited in our analysis (see Supplementary Fig. 4 and 5).

#### SNP filtering

After removing monomorphic SNPs, non-biallelic SNPs, with a *callrate* lower than or equal to 0.99 and with a Hardy-Weinberg disequilibrium test p-value less than or equal to 10^*−*7^, 616 181 autosomal SNPs are kept.

#### CNV filtering

On all our 10 000 individual-subsample, we have 416 970 unique and non-monomorphic positions. When we removed the 2 793 filtered individuals, only 212 054 non-monomorphic positions remain in the CNV count matrix.

### CNV count frequencies

Here, we have computed frequencies on the positions/SNPs that reference CNVs. Then, the same CNV can be referenced by several different SNPs, and a SNP can also reference different CNVs in different individuals (intersecting CNVs). So, here, the frequency of a CNV is the frequency of the presence or absence of CNVs at a given position, for simplicity of writing (see Materials for details). We also use the term common CNV when this frequency is strictly greater than 1% and the term rare CNV when this frequency is less than or equal to 1%.

Among the 212 054 CNVs positions, we have 105 254 positions where only a single individual has a number of copies different from 2 (49.64%). Finally, there are only 1 231 positions with common CNVs, compared with 210,823 with rare CNVs, corresponding to around 0.58% and 99.42% of positions respectively (see Fig. 1).

**Fig. 1.**
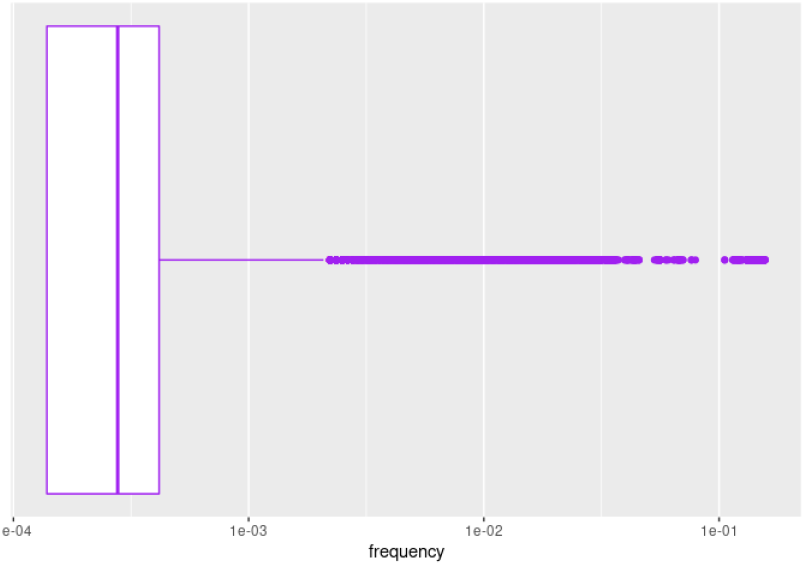
Boxplot of CNV frequencies (count matrix) after data cleaning

### Simulations

After applying linear mixed models on each simulation datasets for each scenario, we can then construct boxplots to study the behavior of our heritability estimates and compare the values obtained with the theoretical value(s) previously set.

#### SNP-dependent phenotype

For this scenario, we have chosen to show results for *τ*_*snp*_ = 3, *σ*^2^ = 7, so 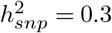 (see Fig. 2).

**Fig. 2.**
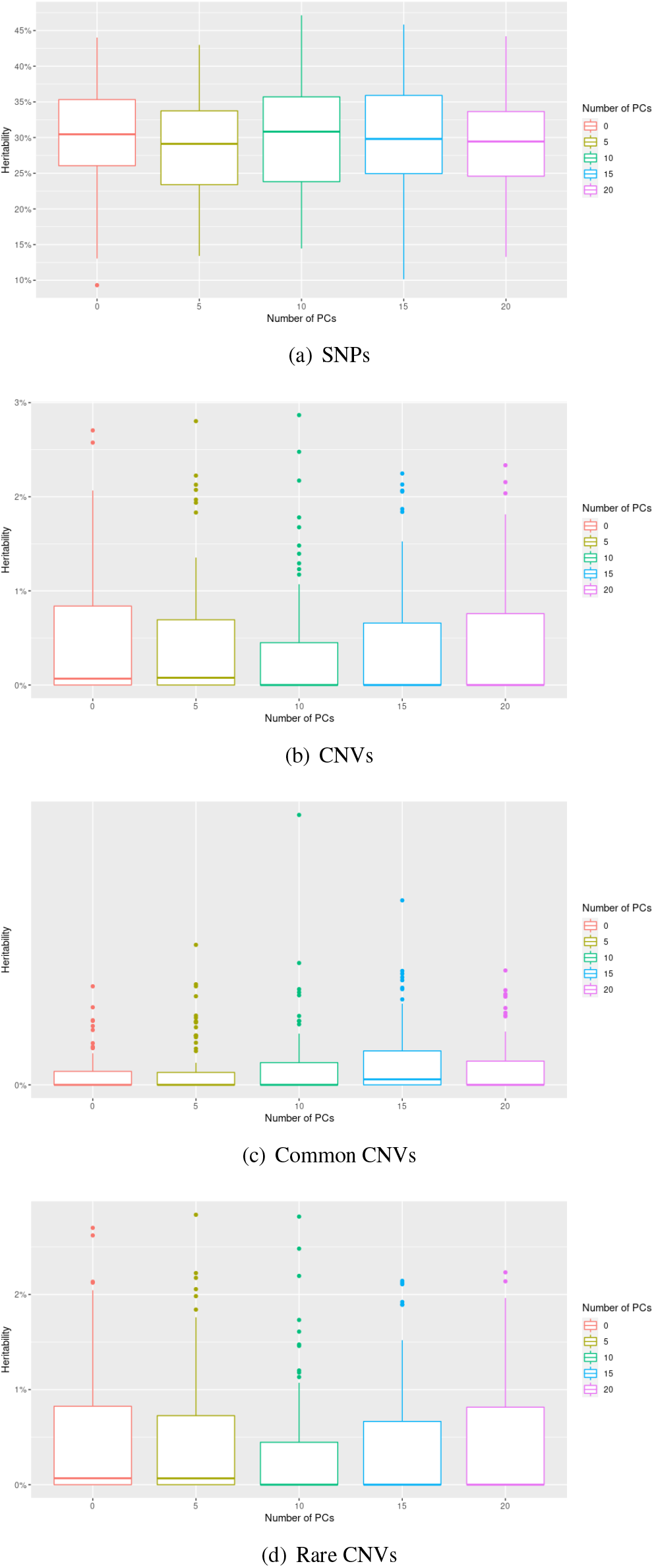
Boxplots of heritability estimated for SNPs, CNVs, common CNVs and rare CNVs, as a function of the number of principal components, when the phenotype depends on SNPs

If we use SNPs in the estimation model (Eq. (7)), i.e. if we use the same parameters as the simulation model, we find that the median of the estimates is generally close to the true heritability (see Fig. 2(a)).

If we use CNVs in the estimation model (Eq. (8)), the estimated heritability is generally close to 0 (see Fig. 2(b)). This is consistent with the fact that our simulated phenotype depends only on SNPs. We note that estimates with common CNVs give lower results than those with rare CNVs, which is certainly due to the fact that we have many more rare CNVs than common ones and the higher standardized values of rare CNVs counts (see Fig. 2(c) and (d)).

#### CNV-dependent phenotype

For the second scebario, we have chosen as parameters *τ*_*cnv*_ = 1, *σ*^2^ = 3, so 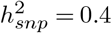 (see Fig. 3).

**Fig. 3.**
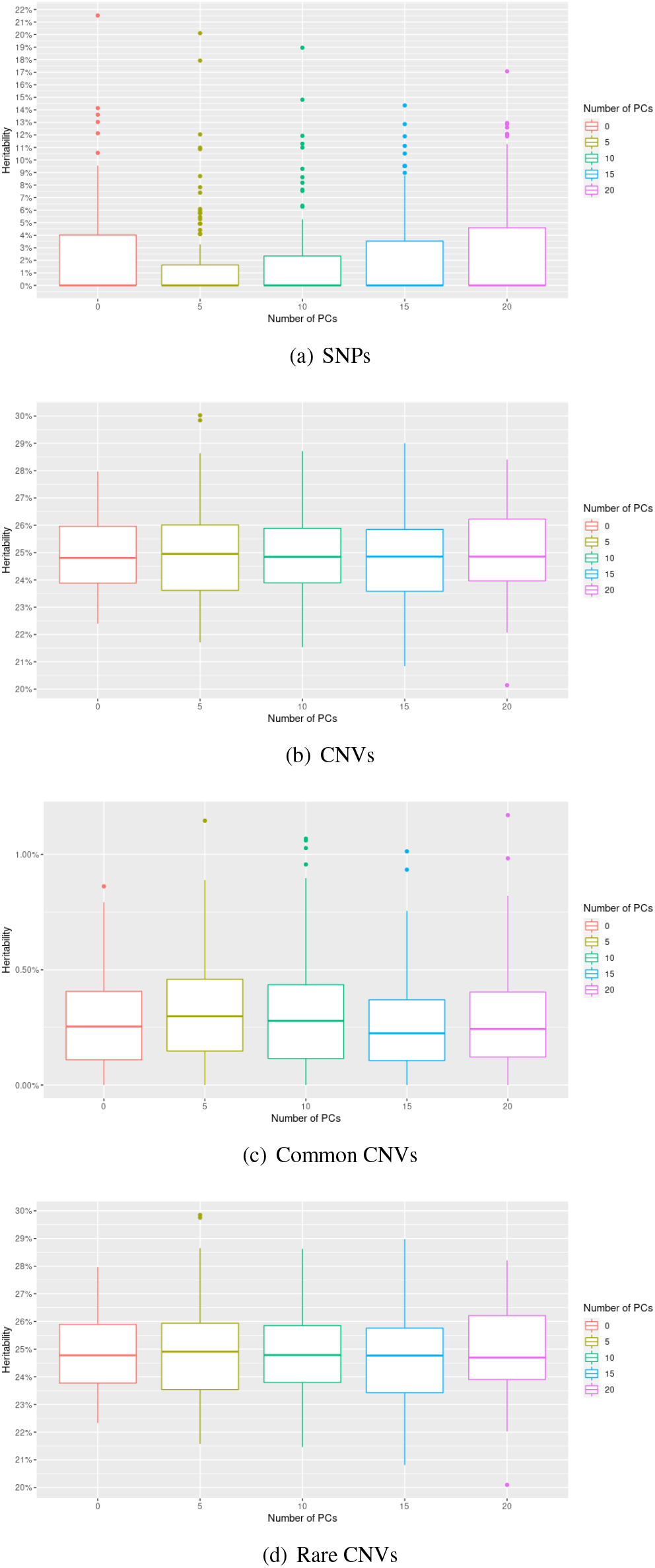
Boxplots of heritability estimated for SNPs, CNVs, common CNVs and rare CNVs, as a function of the number of principal components, when the phenotype depends on CNVs

If we use SNPs in the estimation model (Eq. (7)), the median of the estimates is always equal to 0 (see Fig. 3(a)). This is consistent with the fact that our simulated phenotype depends solely on CNVs. However, we note the presence of extreme values.

If we use the CNVs in the estimation model (Eq. (8)), i.e. if we use the same parameters as the simulation model, we see that the median of the estimates is generally close to the true heritability (see Fig. 3(b)).

Once again, estimates with common CNVs give weaker results than those with rare CNVs (see Fig. 3(c) and (d)).

#### SNP- and CNV-dependent phenotype

Here, we applied in simulations the parameter values *τ*_*snp*_ = 4, *τ*_*cnv*_ = 2, *σ*^2^ = 4, i.e. 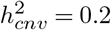 and 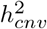 (see Fig. 4).

**Fig. 4.**
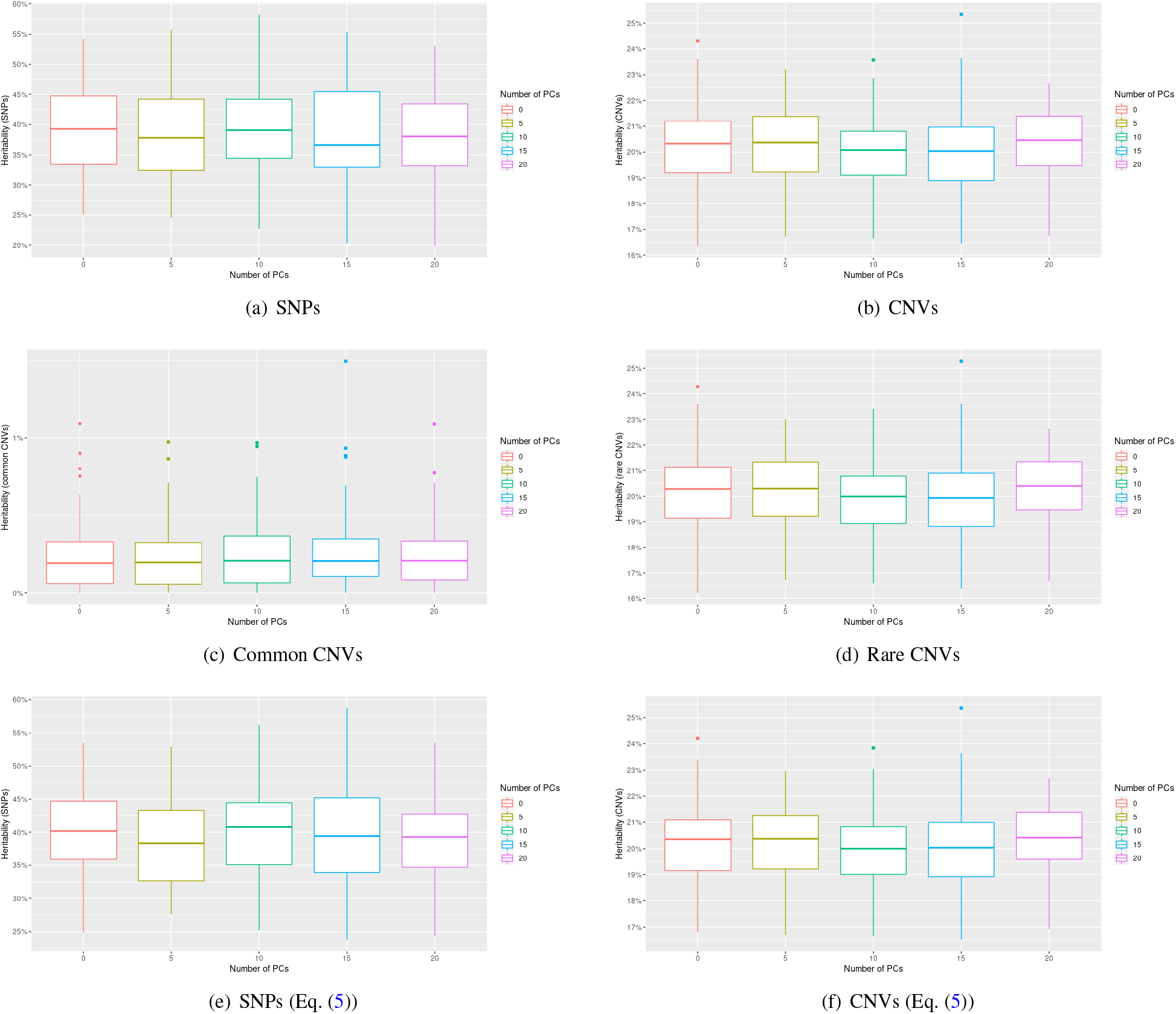
Boxplots of heritability estimated for SNPs, CNVs, common CNVs and rare CNVs, as a function of the number of principal components, when the phenotype depends on SNPs and CNVs

If we use SNPs in the estimation model (Eq. (7)), the median of the estimates is always lower than the true heritability, although it is generally close to the latter (see Fig. 4(a)). Thus, we tend to slightly underestimate the heritability of SNPs.

If we use CNVs in the estimation model (Eq. (8)), we find that the median of the estimates is generally close to the true heritability (see Fig. 4(b)).

As there are very few common CNVs compared to rare CNVs, it is not surprising to see that the results obtained with CNVs are very similar to those obtained with rare CNVs, and that those obtained with common CNVs are all very close to 0 (see Fig. 4(c) and (d)).

If both SNPs and CNVs are used in the estimation model (Eq. (5)), i.e. if we use the same parameters as the simulation model, the results for SNPs are better than those obtained with model defined in Eq. (7), while those for CNVs are very similar to those obtained with model defined in Eq. (8), which is reassuring (see Fig. 4(e) and (f)).

### Heritability estimates on UK Biobank data

#### UKBioBank phenotypes

Here, we performed heritability estimations for SNPs and CNVs on the UKBiobank phenotypes, and more specifically on the 7 207 selected individuals. We were mainly interested in the height of these individuals (see Supplementary Fig. 6).

#### Permuted Phenotypes

Before making final estimates, we first checked that our models were working properly by swapping phenotype values. Specifically, we first permuted the phenotype values which corresponds to phenotype values on the null model. Then, we do not expect to be able to detect an effect of our markers on them, and so we expect the estimated heritability for SNPs or CNVs to be zero, whatever model we use. Finally, we repeated this procedure 100 times for each of the three models. The results are shown in Fig. 5. We first note that the results obtained for SNPs (respectively CNVs) are similar between the different linear mixed models. We can then note that for SNPs, heritability is in at least 50% of cases equal to 0% (median = 0%) as expected, for both models. However, it is important to highlight the fact that the third quartile is around 4% (Eq. (5), see Fig. 5(c)) or 4.3% (Eq. (7), see Fig. 5(a)), and that there are also extreme values for both models reaching around 17%. This may, however, be due to the fact that we come across cases with non-zero heritability in certain permutations (e.g. due to individuals with extreme values for height and with rare variants in common), and not necessarily to an over-estimation problem.

**Fig. 5.**
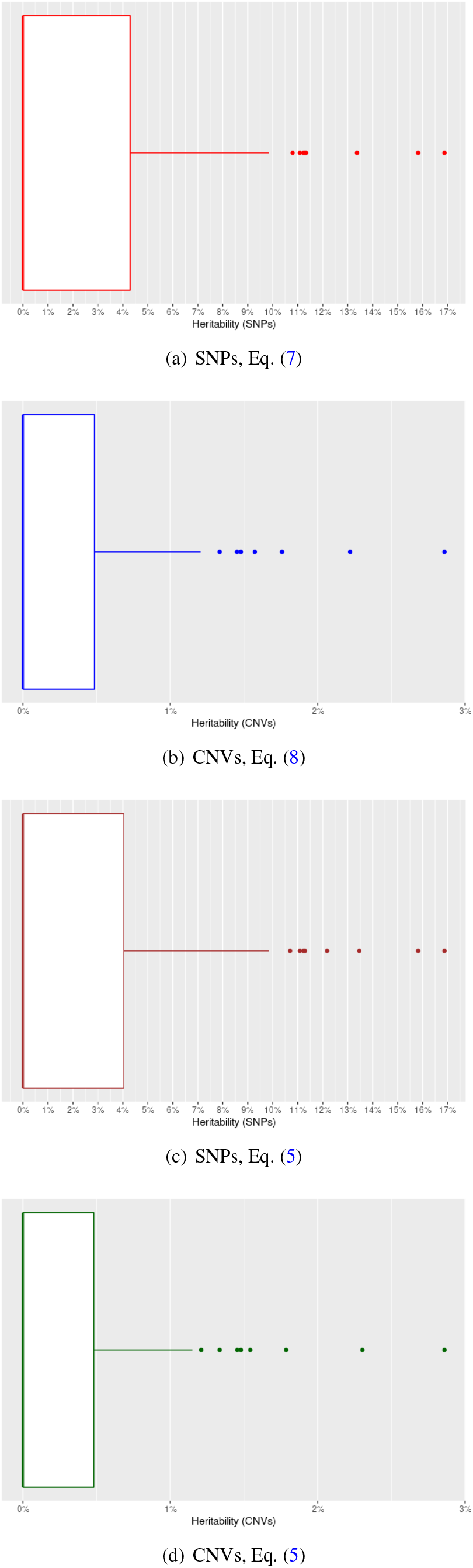
Boxplots of heritability estimated for height (on 100 permutations).

For CNVs, the results are much more stable. Indeed, the median is equal to 0% and the third quartile to around 0.5% for both models (see figures Fig. 5(b) and (d)). In addition, the largest estimated value is less than 3%, again for both models. These results can be explained by the fact that there are probably fewer CNVs profiles that can create heritability in our data, compared with SNPs. Indeed, CNVs have very low frequencies (see Fig. 1). Remember that 2 871 individuals have no CNVs detected. It should also be remembered that of the 212 054 positions with CNVs, there are 105 254 where only one individual has a copy number other than 2 (corresponding to around 49.64% of positions). So we probably have less variability than with SNPs here.

#### Height

Results obtained with models defined in Eq. (7) and Eq. (8) were very similar to those obtained with model in Eq. (5), we thus only show results obtained with the two first models.

Without resampling, estimated heritability for SNPs ranges from around 89.40% to 90.30% across the number of PCs (see Fig. 6(a)). These are higher values than those known for height. Indeed, the heritability of height is estimated at around 80% in the literature (see (9)).

**Fig. 6.**
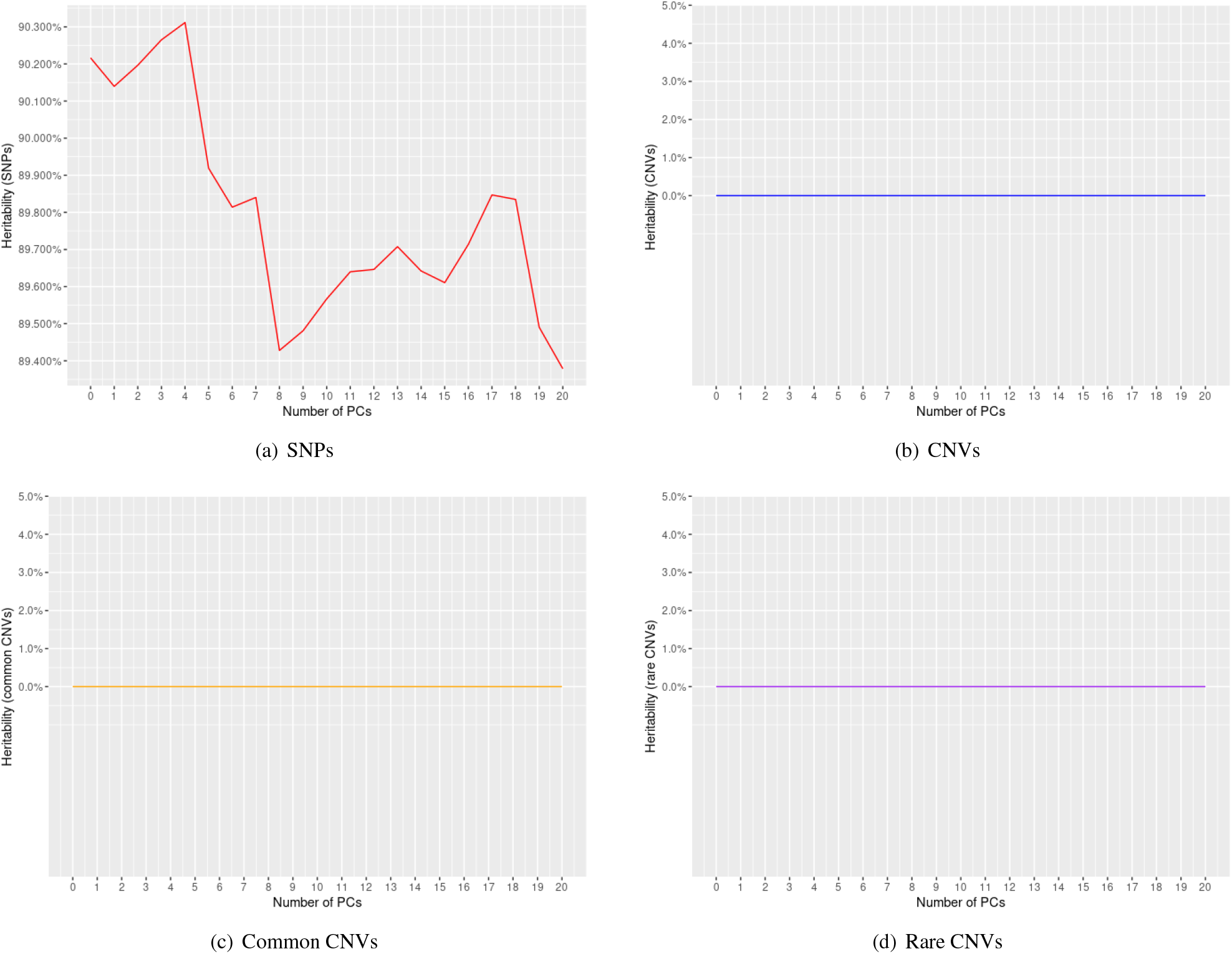
Heritability estimated for height, as a function of the number of principal components.

CNVs have zero heritability (see Fig. 6(b)). The same is true for common and rare CNVs (see Fig. 6(c) and (d)).

With resampling, for SNPs, the median is always between around 88% and 90% (see figure Fig. 7(a)), which is consistent with the estimates in the previous subsection. However, we note the presence of extreme values, such as 100%, and relatively large interquartile ranges, illustrating the variability of our estimates. This variability was also observed on permuted phenotypes.

**Fig. 7.**
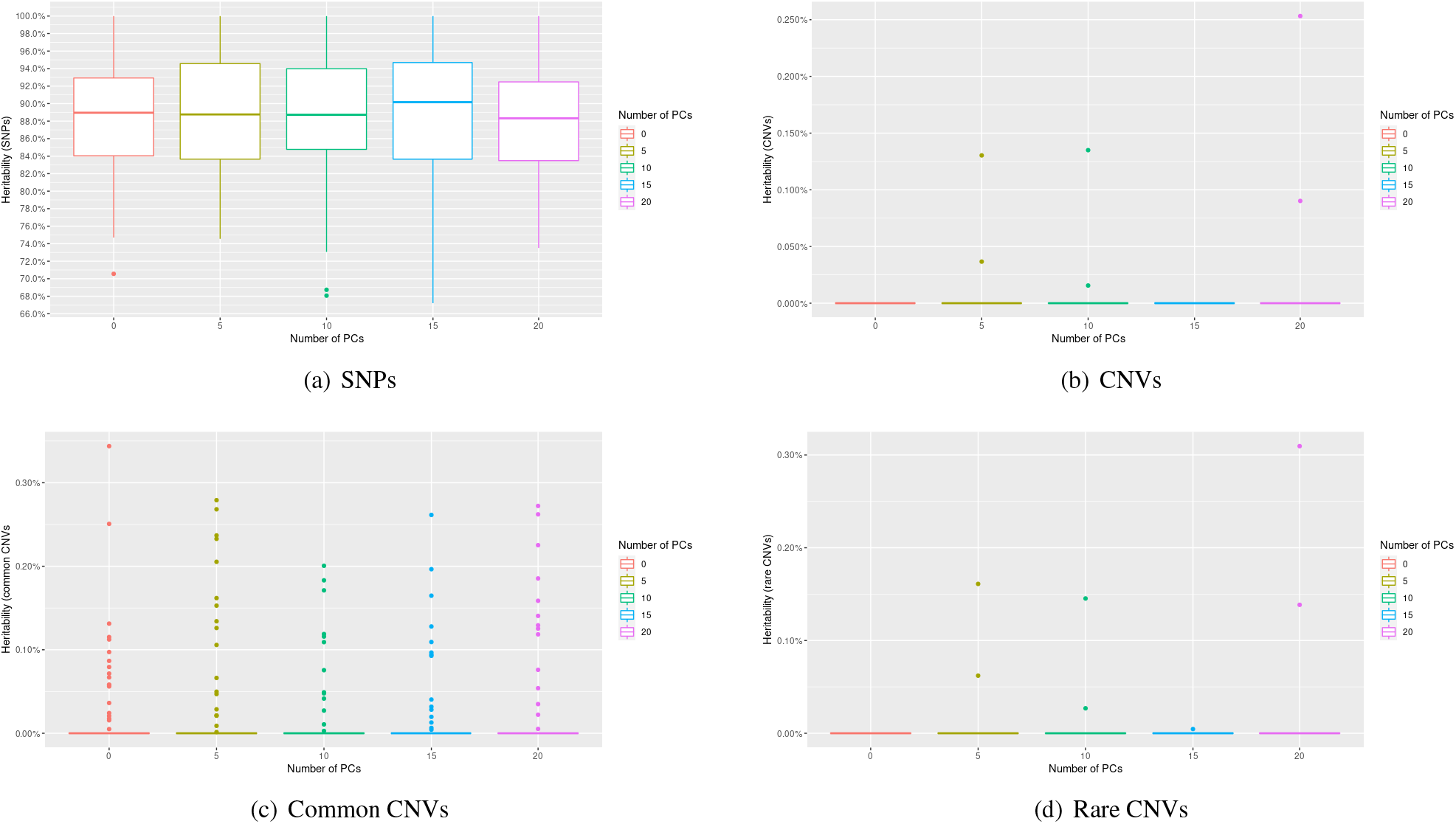
Boxplots of heritability estimated for height, as a function of the number of principal components (resampling of 5000 individuals, without replacement)

For CNVs, estimates are generally equal to 0%, with extreme values below 0.30% (see Fig. 7(b)), which again is not surprising according to the estimates in the previous subsection. Thus, estimates for CNVs are very stable here, which is consistent with the results of simulations where we already observed low variability in estimates.

The observation is similar if we retain only common or rare CNVs (see Fig. 7(c) and (d)).

#### Hypertension

We studied one disease (binary phenotype), the hypertension which is the most common in UKBioBank dataset (22.69%). As with heritability estimates for height, estimates without and with resampling have been carried out here. The results obtained without resampling are summarized in figure Fig. 8, and those with resampling in figure Fig. 9.

**Fig. 8.**
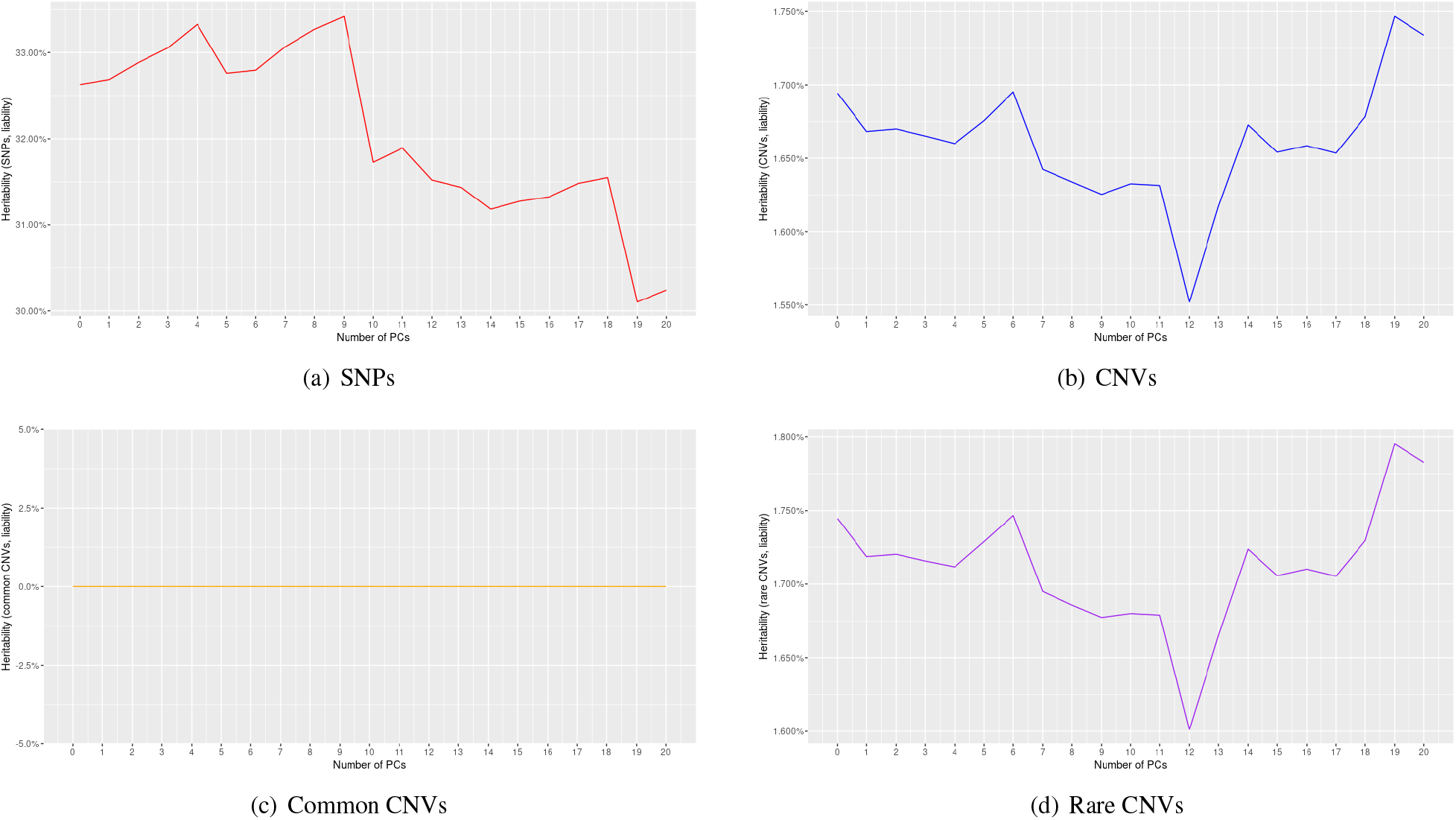
Heritability estimated for hypertension, under the liability scale, as a function of the number of principal components

**Fig. 9.**
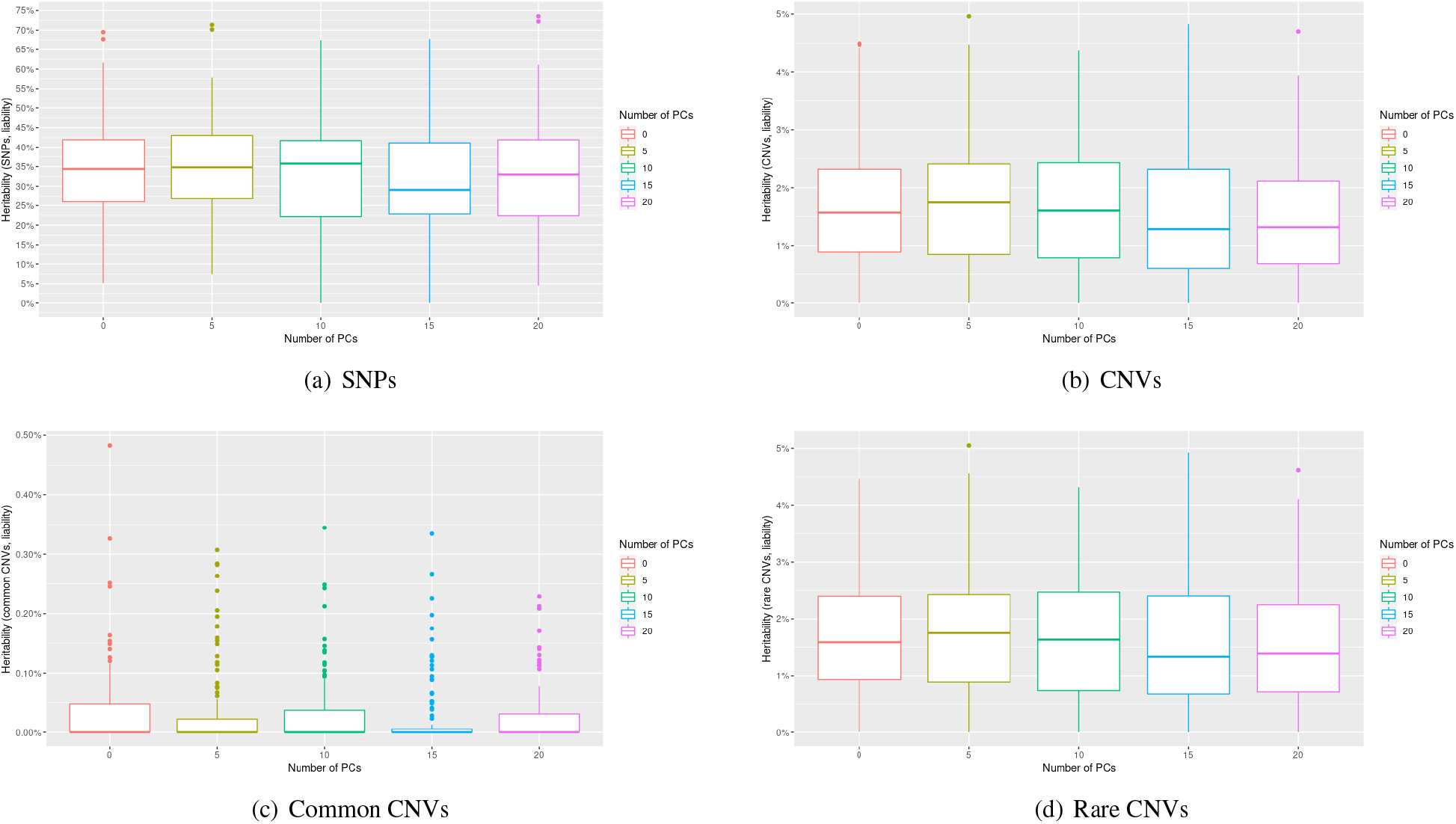
Boxplots of heritability estimated for hypertension, under the liability scale, as a function of the number of principal components (resampling of 5000 individuals, without replacement).

Concerning the results without resampling, we note that CNVs do not have zero heritability here, although it remains very low (it oscillates between around 1.55% and 1.75%). On all CNVs. For SNPs, heritability oscillates between around 30% and 33.5%. These are much lower values than those obtained for height, which can be explained by the fact that this phenotype is less heritable.

Regarding the results with resampling, we again note the instability of SNP heritability estimates. However, we note that the medians are always close to the values obtained without resampling, which is reassuring. For CNVs, the results are much more stable, and medians are also close to the values estimated without resampling (the observation is similar for common or rare CNVs).

## Discussion

### Simulation analysis

First, we note the variability of our estimates. Indeed, the interquartile ranges, whether for estimates using SNPs or CNVs, are large. Moreover, including principal components in the estimation model does not seem to have much influence on the estimates (the boxplots are all relatively similar), which is expected given that we do not include principal components when simulating the phenotype. It is also important to note that in simulation scenarios (1), (2), (3) and (4) estimating with the model defined in Eq. (5), or using the Eq. (7) and Eq. (8) models separately, gives similar results (results not shown), which is reassuring because it shows that in the Eq. (5) model, we are not capturing the effect of CNVs with SNPs or vice versa, and this corresponds well to what is expected (we are not taking interactions into account in the model).

Other simulations were also carried out (results not shown) by including the first 10 principal components as features. We remarked that all the results obtained are very similar between the two approaches, which may be explained by the fact that our population is homogeneous or that the effect we applied to the 10 principal components via the choice of *β ∼ N* (0_*R*_15, 1) is not strong enough.

### Real Phenotype, the height and hypertension

We can formulate several possibilities to explain our results on height phenotype:

1. CNV data modeling problems;

2. convergence problem of the linear mixed model (for CNVs);

3. CNV information already taken into account by SNPs;

4. CNV effects too weak to be detected;

5. no effect of CNVs;

6. technology problem (genotyping chips and/or CNV detection pipeline).

If the first possibility were true, then we would not be able to find the true heritabilities in the simulations data.

To investigate the second possibility, we looked at the likelihood of model defined by Eq. (8), without including fixedeffects principal components. Specifically, we displayed the log-likelihood level curves as a function of the values of *τ*_*cnv*_ and *σ*^2^ (see Supplementary Fig. 7(a)), as well as the likelihood as a function of the heritability of the CNVs 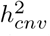 (see Supplementary Fig. 7(b)). We can then see that there is no convergence problem. Indeed, the model does converge, but it converges for a negative value of *τ*_*cnv*_ (and therefore of). Now, we are looking for the maximum likelihood under the constraint that *τ*_*cnv*_ *≥* 0, hence 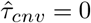 (and therefore 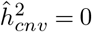)

As for the third possibility, this is highly unlikely, given that the one random linear mixed models give very similar estimates to those of the complete model.

In the end, only the last three possibilities seem relevant. In order to determine which of the three is the most plausible, we need to gain in power. One possible approach is to form groups of CNVs (called CNV Regions, abbreviate to CNVRs), rather than working on each CNV individually.

This is the approach used by the CCRET method (13)) where two CNVs belong to the same region as soon as they have a base pair in common. This method also makes it possible to evaluate the effect of CNV length and their intersection with known genes (which we have not taken into account in our calculations), in addition to the effect of their dosage (i.e. copy number). We wanted to apply the CCRET method to our 7 207 individuals, but ran up against a problem of calculation time (more than a week to calculate the three matrices). To solve this problem, we restricted ourselves to calculating the CNV dosage matrix, which is sufficient for heritability estimates. In fact, all we have to do is use it instead of *K*_*cnv*_ in our models. In doing so, we again obtain an estimated heritability of zero.

We think that the last two possibilities are the most plausible, i.e. an absence of CNV effect for this phenotype o a problem linked to the technology used to extract the CNVs. With regard to the technology problem in particular, we note that the vast majority of CNVs extracted are single-copy deletions, reflecting the lack of diversity in our CNV data (see Supplementary Fig. 8). This may be due to a CNV detection problem (i.e. the chips used may not be suited to this task), but we cannot verify this.

## ACKNOWLEDGEMENTS

This research has been conducted using the UK Biobank Resource under Application Number 45408.

## Notes

### Competing Interest Statement

The authors have declared no competing interest.

## Bibliography

1. Matthew Aguirre, Manuel A. Rivas, and James Priest. Phenome-wide burden of copynumber variation in the UK biobank. The American Journal of Human Genetics, 105(2): 373–383, August 2019. doi: 10.1016/j.ajhg.2019.07.001.

2. David Owen, Mathew Bracher-Smith, Kimberley M. Kendall, Elliott Rees, Mark Einon, Valentina Escott-Price, Michael J. Owen Michael C. O’Donovan, and George Kirov. Effects of pathogenic CNVs on physical traits in participants of the UK biobank. BMC Genomics, 19(1), December 2018. doi: 10.1186/s12864-018-5292-7.

3. Aurélien Macé, Marcus A. Tuke, et al. CNV-association meta-analysis in 191, 161 european adults reveals new loci associated with anthropometric traits. Nature Communications, 8(1), September 2017. doi: 10.1038/s41467-017-00556-x.

4. Chiara Auwerx, Maarja Lepamets, Marie C Sadler, Marion Patxot, Miloš Stojanov, David Baud, Reedik Mägi, Tõnu Esko, Andres Metspalu, Lili Milani, et al. The individual and global impact of copy-number variants on complex human traits. The American Journal of Human Genetics, 109(4):647–668, 2022.

5. Cathie Sudlow, John Gallacher, Naomi Allen, Valerie Beral, Paul Burton, John Danesh, Paul Downey, Paul Elliott, Jane Green, Martin Landray, Bette Liu, Paul Matthews, Giok Ong, Jill Pell, Alan Silman, Alan Young, Tim Sprosen, Tim Peakman, and Rory Collins. Uk biobank: An open access resource for identifying the causes of a wide range of complex diseases of middle and old age. PLOS Medicine, 12(3):1–10, 03 2015. doi: 10.1371/journal.pmed.1001779.

6. Clare Bycroft, Colin Freeman, Desislava Petkova, Gavin Band, Lloyd T Elliott, Kevin Sharp, Allan Motyer, Damjan Vukcevic, Olivier Delaneau, Jared O’Connell, et al. Genome-wide genetic data on^∼^ 500,000 uk biobank participants. BioRxiv, page 166298, 2017.

7. Jian Yang, Teri A Manolio, Louis R Pasquale, Eric Boerwinkle, Neil Caporaso, Julie M Cunningham, Mariza de Andrade, Bjarke Feenstra, Eleanor Feingold, M Geoffrey Hayes, William G Hill, Maria Teresa Landi, Alvaro Alonso, Guillaume Lettre, Peng Lin, Hua Ling, William Lowe, Rasika A Mathias, Mads Melbye, Elizabeth Pugh, Marilyn C Cornelis, Bruce S Weir, Michael E Goddard, and Peter M Visscher. Genome partitioning of genetic variation for complex traits using common SNPs. Nature Genetics, 43(6):519–525, May 2011. doi: 10.1038/ng.823.

8. K. Wang, M. Li, D. Hadley, R. Liu, J. Glessner, S. F.A. Grant, H. Hakonarson, and M. Bucan. PennCNV: An integrated hidden markov model designed for high-resolution copy number variation detection in whole-genome SNP genotyping data. Genome Research, 17(11): 1665–1674, November 2007. doi: 10.1101/gr.6861907.

9. Peter M. Visscher, William G. Hill, and Naomi R. Wray. Heritability in the genomics era — concepts and misconceptions. Nature Reviews Genetics, 9(4):255–266, March 2008. doi: 10.1038/nrg2322.

10. Arthur R Gilmour, Robin Thompson, and Brian R Cullis. Average information reml: an efficient algorithm for variance parameter estimation in linear mixed models. Biometrics, pages 1440–1450, 1995.

11. Sang Hong Lee, Naomi R. Wray, Michael E. Goddard, and Peter M. Visscher. Estimating missing heritability for disease from genome-wide association studies. The American Journal of Human Genetics, 88(3):294–305, March 2011. doi: 10.1016/j.ajhg.2011.02.002.

12. 46th European Mathematical Genetics Meeting (EMGM) 2018, Cagliari, Italy, April 18-20, 2018: Abstracts. Human Heredity, 83(1):1–29, 04 2018. ISSN 0001-5652. doi: 10.1159/000488519.

13. Jung-Ying Tzeng, Patrik K. E. Magnusson, Patrick F. Sullivan, and Jin P. Szatkiewicz and. A new method for detecting associations with rare copy-number variants. PLOS Genetics, 11(10):e1005403, October 2015. doi: 10.1371/journal.pgen.1005403.

