## Supplementary Materials for "Copy Number Variants and heritability estimates on UKBiobank data"

### Supplementary Note 1: CNVs extraction

In addition to SNPs, CNVs were extracted from SNP data using the **PennCNV** tool (1) and the pipeline described in (2). During this extraction, 2 871 individuals were excluded, as no CNVs were detected for them. Once the extraction is complete, we obtain various information for each of the CNVs.

- the chromosome where the CNV is located;
- the start and end position of the CNV;
- the length of the CNV;
- the number of copies of the CNV and the *copy number state*;
- the number of SNPs marking the CNV;
- the first and last SNPs marking the CNV;
- the identifier of the individual possessing this CNV.

Then, we computed the exact number of CNVs detected for each individual.
Referring to the article (1), the link between *copy number state* and the number of copies of a CNV can be summarized in the following table:

| State | Total copy number | Description (autosome) |
|---|---|---|
| 1 | 0 | Deletion of two copies |
| 2 | 1 | Deletion of one copy |
| 3 | 2 | Normal state |
| 4 | 2 | Copy-neutral with LOH |
| 5 | 3 | Single copy duplication |
| 6 | 4 | Double copy duplication |

The term *LOH* stands for *Loss of Heterozygosity*. In other words, Copy-neutral with LOH refers to loss of heterozygosity with no change in copy number. If we note AA, AB and BB as the genotypes for any SNP, this means that a SNP with a *copy number state* equal to 4 has either the AA or BB genotype, but never the AB genotype. Thus, the heterozygous genotype is lost, but the copy number is normal (two alleles are present).
From the raw data, we can build a second dataset, this time containing information about the SNPs referencing the CNVs. More precisely, what we call **CNV count matrix** in the following, contains (in columns):

- the chromosome where the SNP is located;
- the position of the SNP;
- the SNP identifier;
- the number of copies of the SNP for each individual (one column per individual).

There are no *copy number state* equal to 4 in our raw CNV data, so the 2 present in this count matrix indicate a normal copy number for the SNP in question and not a loss of heterozygosity with no change in copy number (i.e. there is no ambiguity).
Individuals with no detected CNVs (2 871) were manually added to the CNV count matrix with normal copy number for the all SNPs.

DRAFT

### Supplementary Note 2: Others simulation scenarios

**A. Phenotypes dependent on common CNVs.** We have chosen as parameters for simulations, $\tau_{common\,cnv} = 2$, $\sigma^2 = 18$, so $h^2_{common\,cnv} = 0.1$ (see Supplementary Fig. 9).
If we use SNPs in the estimation model (Eq. (7)), we tend to overestimate the heritability of SNPs (see Supplementary Fig. 9(a)). Indeed, the medians are different from 0 in all cases, and we note the presence of extreme values. Note that when we include 5 principal components, the estimates are closer to those expected (the median is close to $0\%$ and the estimates are less scattered). These results are unexpected given that the simulated phenotype depends only on common CNVs.
If we use the CNVs in the estimation model (Eq. (8)), we notice that the estimated heritability is generally lower than that of the common CNVs (see Supplementary Fig. 9(b)). Indeed, the medians are always below $0.1$. This is probably due to the fact that only common CNVs influence phenotype. Taking all CNVs into account when the vast majority are not involved in phenotype simulation probably lowers heritability.

If we use the common CNVs in the estimation model (Eq. (9)), i.e. if we use the same parameters as the simulation model, we find that the median of the estimates is generally close to the true heritability (see Supplementary Fig. 9(c)). With rare CNVs (Eq. (10)), the results are weaker (see Supplementary Fig. 9(d)), but we expected to have estimates generally equal to 0 given that the phenotype does not depend on them.

**B. Phenotypes dependent on rare CNVs.** We have chosen for this scenario $\tau_{rare\,cnv} = 1.5$, $\sigma^2 = 8.5$, so $h^2_{rare\,cnv} = 0.15$ (see Supplementary Fig. 10).

If we use SNPs in the estimation model (Eq. (7)), the median of the estimates is always equal to 0 (see Supplementary Fig. 10(a)). This is consistent with the fact that our simulated phenotype depends only on rare CNVs. However, we note the presence of extreme values.

If we use the CNVs in the estimation model (Eq. (8)), we find that the median of the estimates is generally close to the true heritability (see Supplementary Fig. 10(b)).

As there are very few common CNVs compared to rare CNVs, it is not surprising to see that the results obtained with CNVs are very similar to those obtained with rare CNVs, and that those obtained with common CNVs are all very close to 0 (see Supplementary Fig. 10(c) and (d)).

**C. Phenotype dependent on SNPs and common CNVs.** We have chosen: $\tau_{snp} = 4$, $\tau_{common\,cnv} = 2$, $\sigma^2 = 4$, i.e. $h^2_{snp} = 0.4$ and $h^2_{common\,cnv} = 0.2$ (see Supplementary Fig. 11).

If SNPs are used in the estimation model (Eq. (7)), the median of the estimates is always higher than the true heritability, although it is generally close to the latter (see Supplementary Fig. 11(a)). Thus, we tend to slightly overestimate the heritability of SNPs. The finding is therefore similar to that in simulations with phenotypes dependent on common CNVs.

If we use CNVs, common CNVs or rare CNVs in the estimation model (Eq. (8), Eq. (9) and Eq. (10)), the finding is also similar to that in simulations with phenotypes dependent on common CNVs (see Supplementary Fig. 11(b), (c) and (d)).

If both SNPs and common CNVs are used in the estimation model (Eq. (5)), i.e. if we use the same parameters as the simulation model, the results for SNPs are better than those obtained with Eq. (7), while those for common CNVs are very similar to those obtained with mEq. (8), Eq. (9) and Eq. (10), which is reassuring (see Supplementary Fig. 11(e) and (f)).

**D. Phenotype dependent on SNPs and rare CNVs.** We chose parameters $\tau_{snp} = 4$, $\tau_{rare\,cnv} = 2$, $\sigma^2 = 4$, i.e. $h^2_{snp} = 0.4$ and $h^2_{rare\,cnv} = 0.2$ (see Supplementary Fig. 12.

If we use SNPs in the estimation model (Eq. (7)), we tend to slightly underestimate the heritability of SNPs (see Supplementary Fig. 12(a)).

If we use CNVs, common CNVs or rare CNVs in the estimation model (Eq. (8)), the finding is also similar to that in simulations with phenotypes dependent on rare CNVs (see Supplementary Fig. 12(b), (c) and (d)).

If both SNPs and rare CNVs are used in the estimation model (Eq. (5)), i.e. if we use the same parameters as the simulation model, the results for SNPs are better than those obtained with Eq. (7), while those for rare CNVs are very similar to those obtained with model in Eq. (8),, which is reassuring (see Supplementary Fig. 12(e) and (f)).

### Bibliography


1. K. Wang, M. Li, D. Hadley, R. Liu, J. Glessner, S. F.A. Grant, H. Hakonarson, and M. Bucan. PennCNV: An integrated hidden markov model designed for high-resolution copy number variation detection in whole-genome SNP genotyping data. *Genome Research*, 17(11): 1665–1674, November 2007. doi: 10.1101/gr.6861907.
2. Aurélien Macé, Marcus A. Tuke, et al. CNV-association meta-analysis in 191, 161 european adults reveals new loci associated with anthropometric traits. *Nature Communications*, 8(1), September 2017. doi: 10.1038/s41467-017-00556-x.


DRAFT

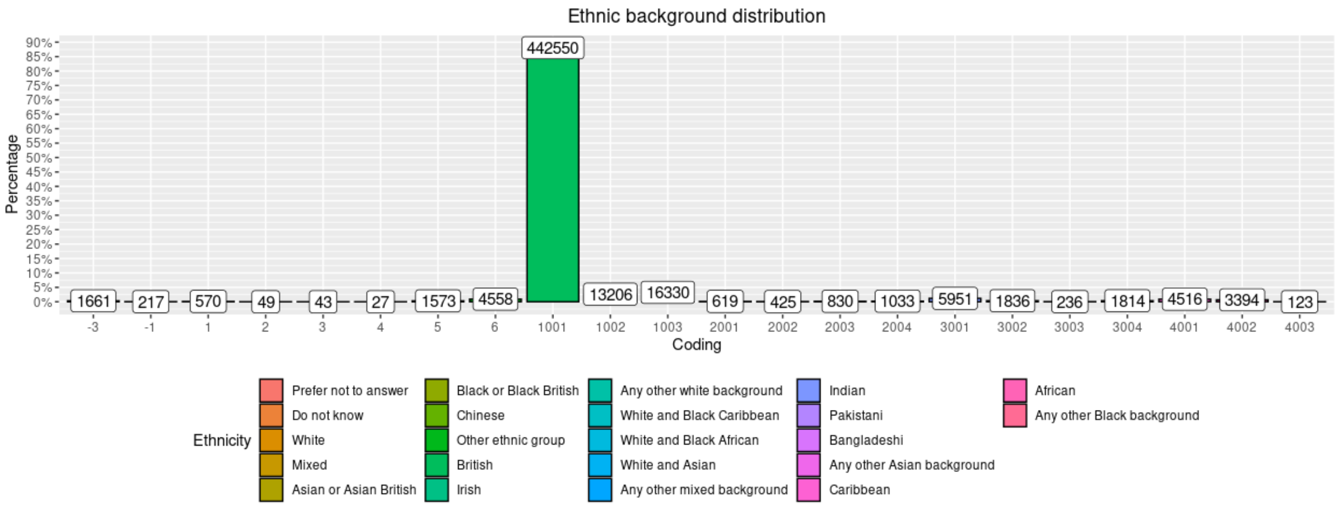


**Supplementary Figure 1.** Individual Ethnic distribution.

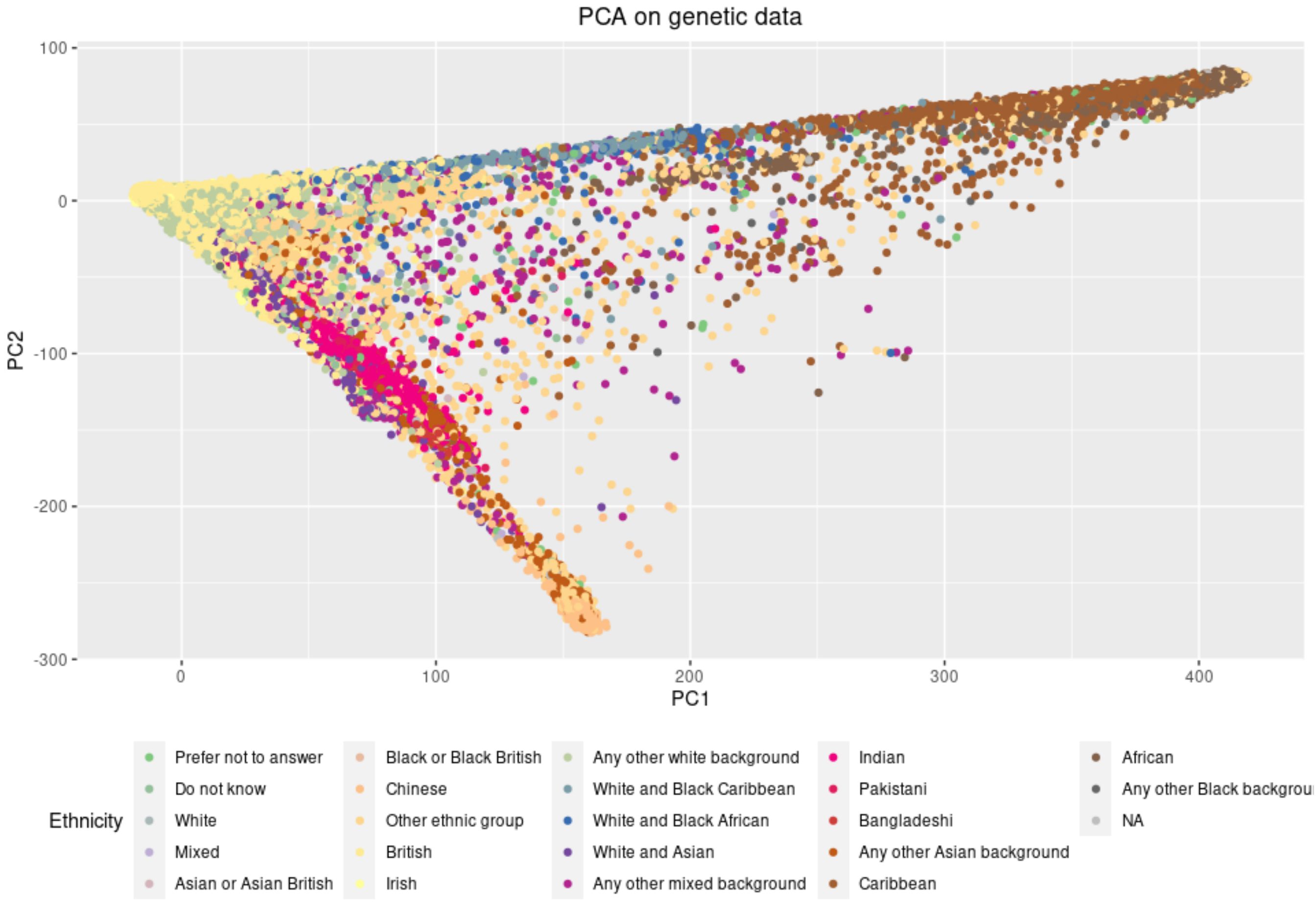


**Supplementary Figure 2.** PCA scores by individual origin, for the first two principal components.

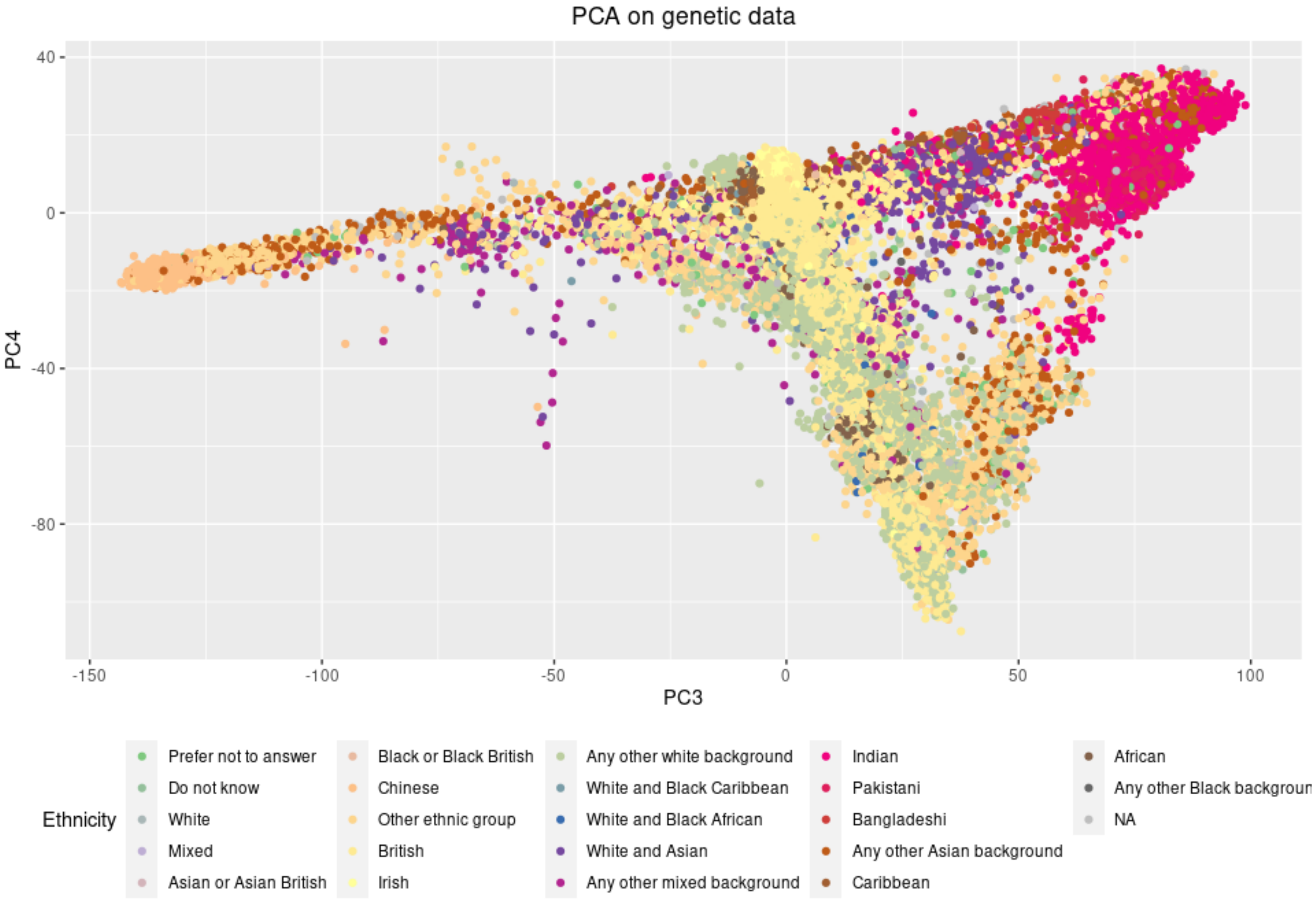


**Supplementary Figure 3.** PCA scores by individual origin, for principal components 3 and 4.

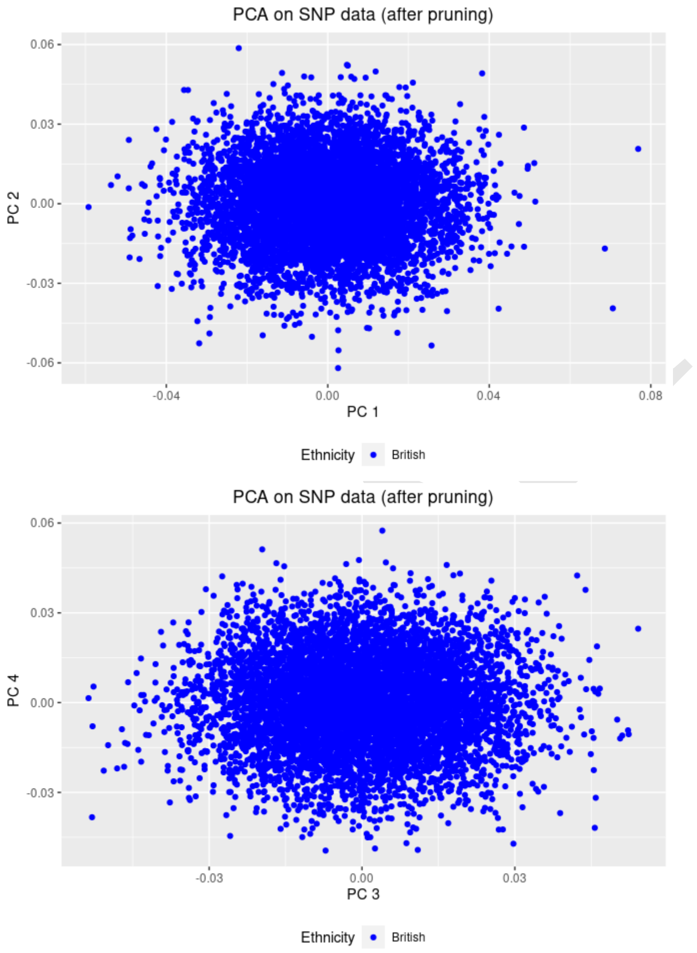


**Supplementary Figure 4.** PCA scores (with GRM after pruning) for components 1 and 2, then 3 and 4

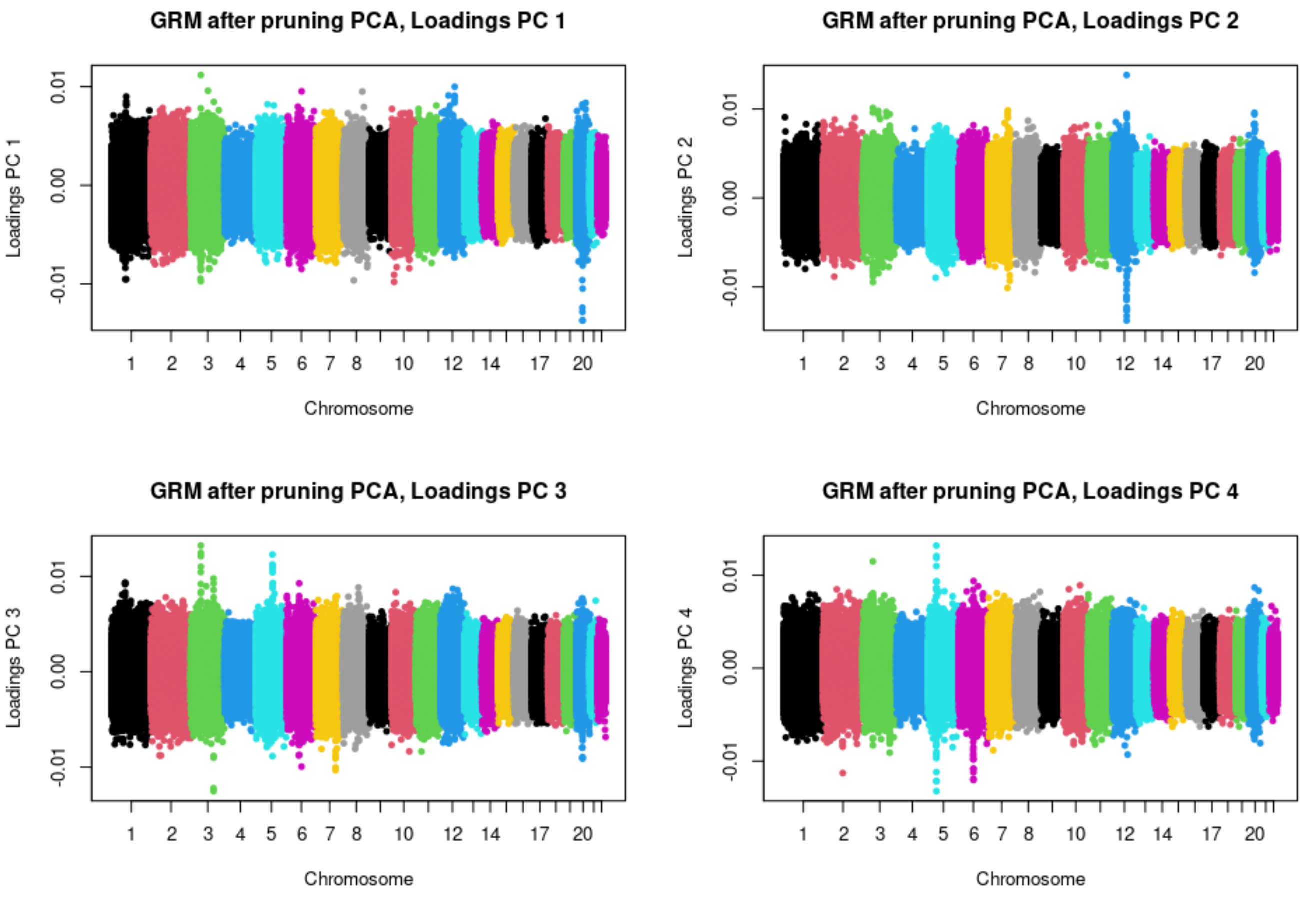


**Supplementary Figure 5.** Loadings for the first four principal components (GRM after pruning)

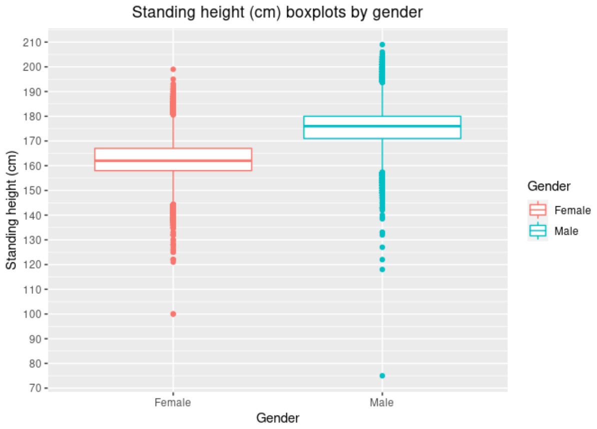


**Supplementary Figure 6.** Height boxplot by sex.

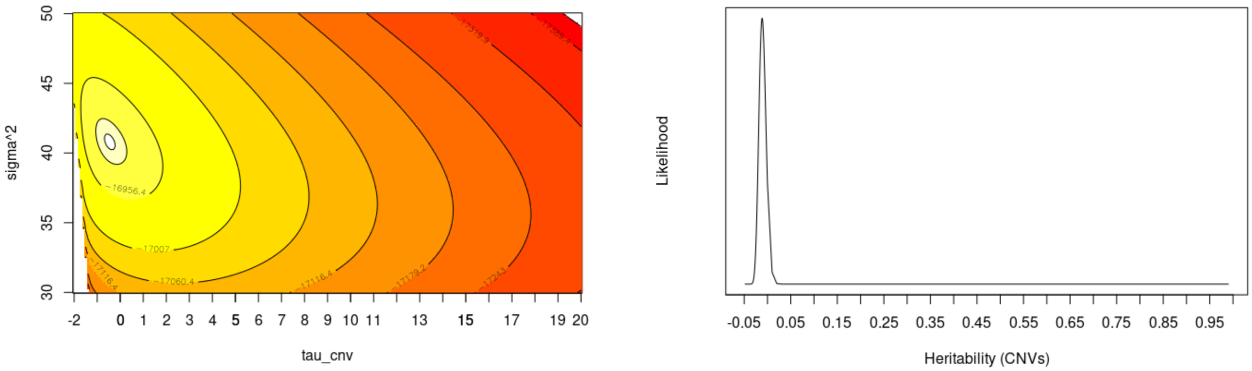


(a) Level curves of the log-likelihood as a function of $\tau_{cnv}$ and $\sigma^2$

(b) Likelihood as a function of CNV heritability $h^2_{cnv}$

**Supplementary Figure 7.** Level curves of the log-likelihood and Likelihood as a function of CNVs variance parameters for height.

AFT

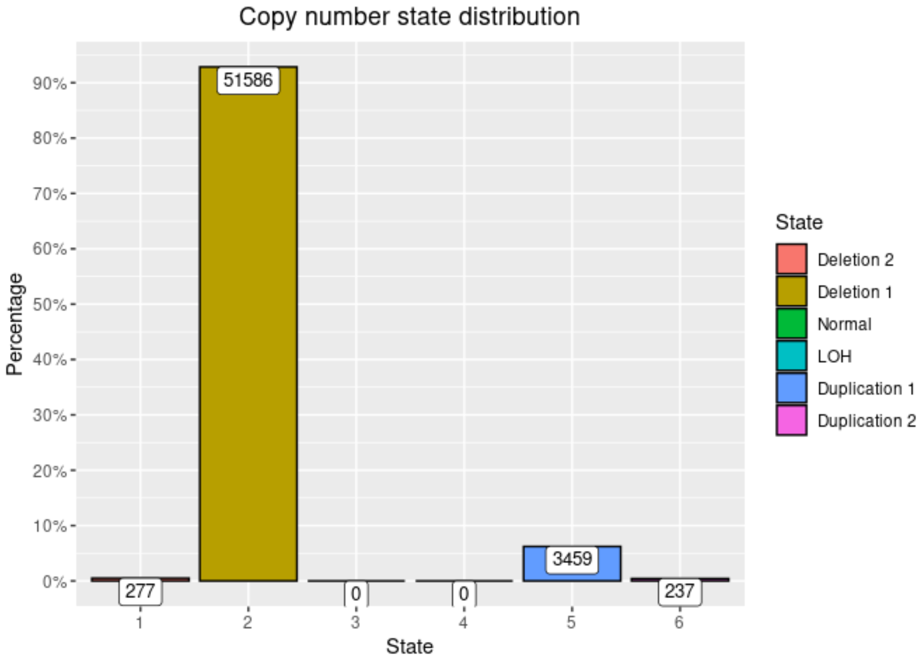


**Supplementary Figure 8.** Distribution of CNV types (raw data)

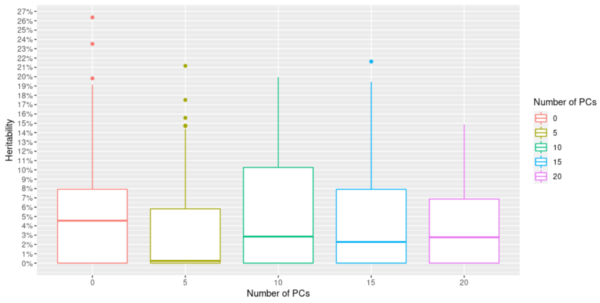


(a) SNPs

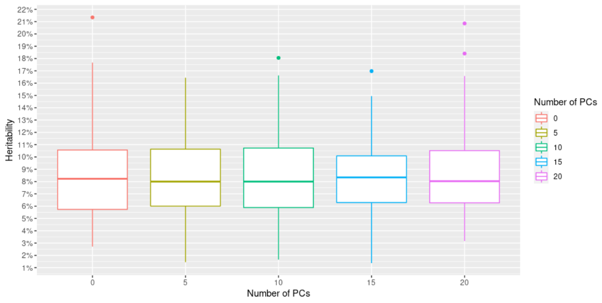


(b) CNVs

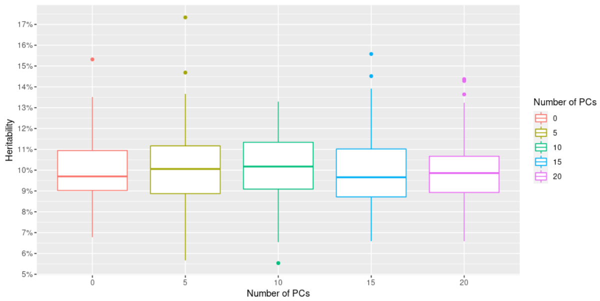


(c) Common CNVs

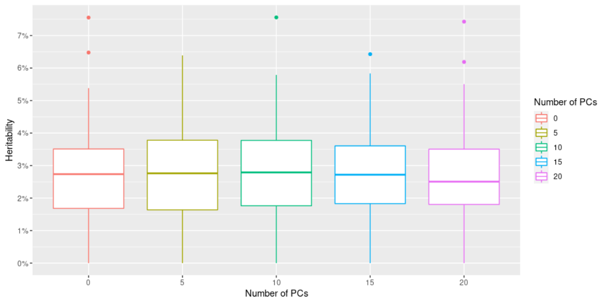


(d) Rare CNVs

**Supplementary Figure 9.** Boxplots of heritability estimated for SNPs, CNVs, common CNVs and rare CNVs, as a function of the number of principal components, when the phenotype depends on common CNVs

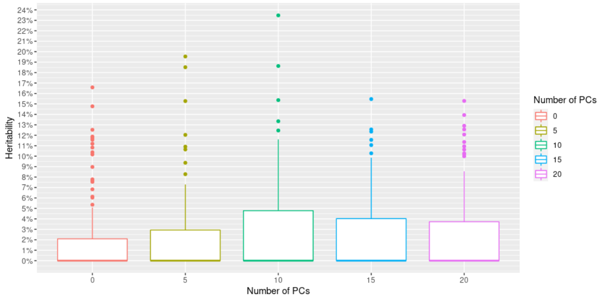


(a) SNPs

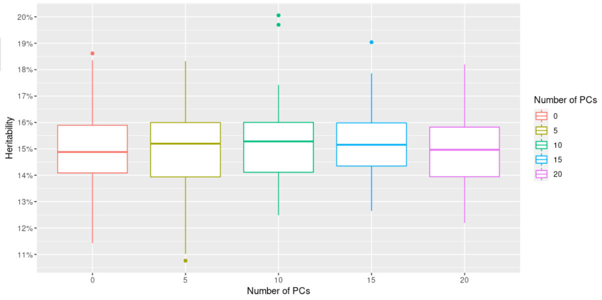


(b) CNVs

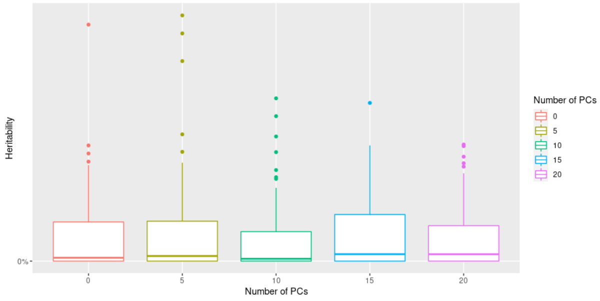


(c) Common CNVs

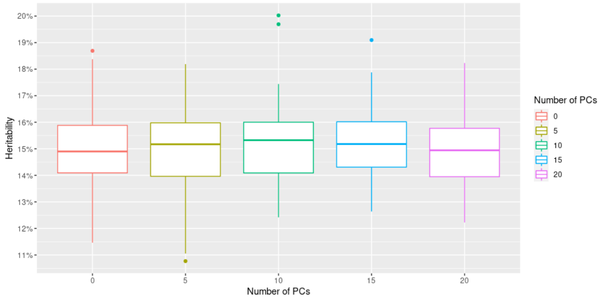


(d) Rare CNVs

**Supplementary Figure 10.** Boxplots of heritability estimated for SNPs, CNVs, common CNVs and rare CNVs, as a function of the number of principal components, when the phenotype depends on rare CNVs

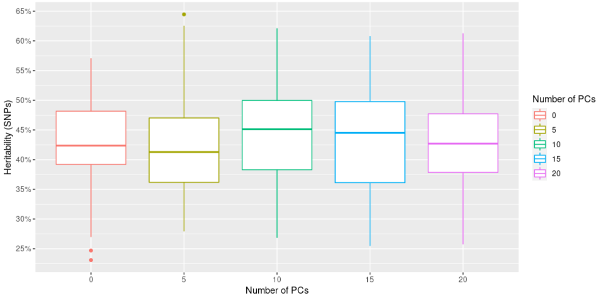


(a) SNPs

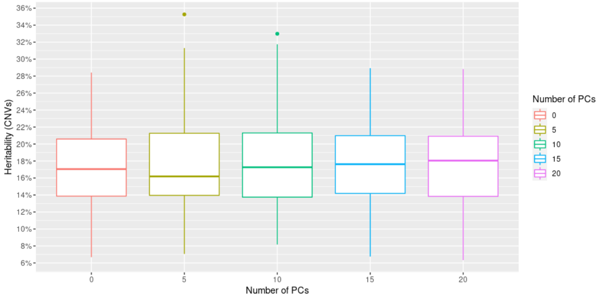


(b) CNVs

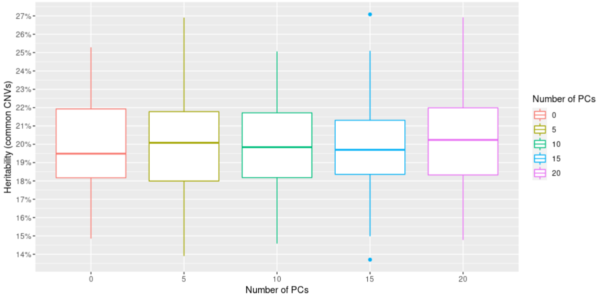


(c) Common CNVs

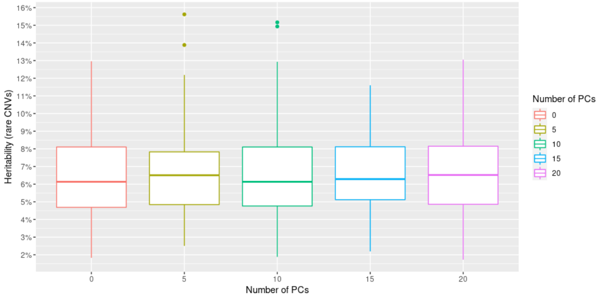


(d) Rare CNVs

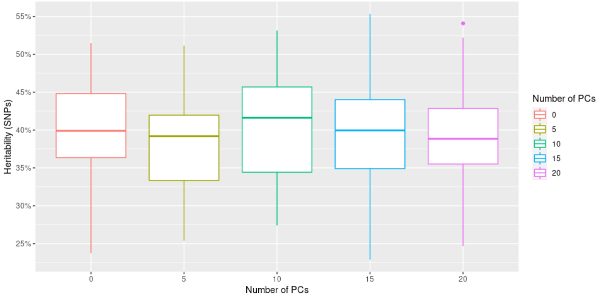


(e) SNPs (Eq. (5))

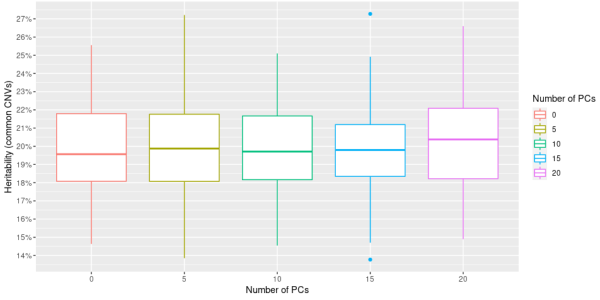


(f) Common CNVs (Eq. (5))

**Supplementary Figure 11.** Boxplots of heritability estimated for SNPs, CNVs, common CNVs and rare CNVs, as a function of the number of principal components, when the phenotype depends on SNPs and common CNVs

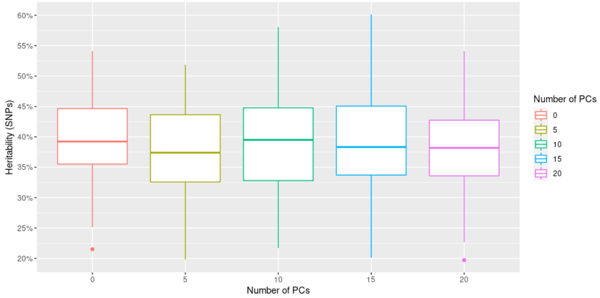


(a) SNPs

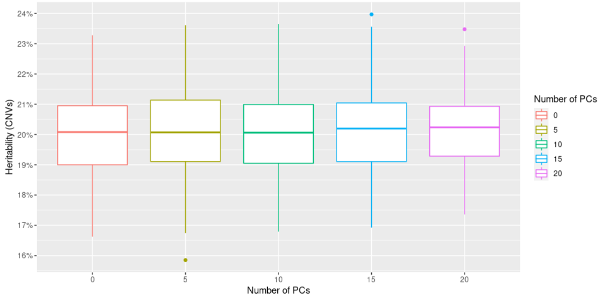


(b) CNVs

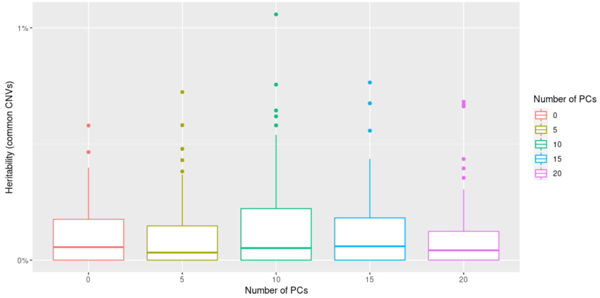


(c) Common CNVs

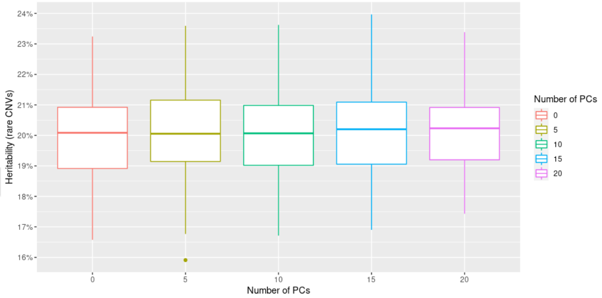


(d) Rare CNVs

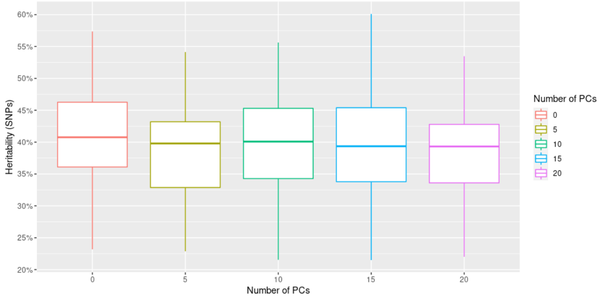


(e) SNPs (Eq. (5))

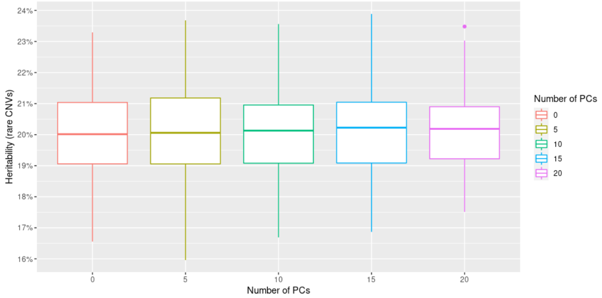


(f) Rare CNVs (Eq. (5))

**Supplementary Figure 12.** Boxplots of heritability estimated for SNPs, CNVs, common CNVs and rare CNVs, as a function of the number of principal components, when the phenotype depends on SNPs and rare CNVs
